# Cuticular hydrocarbon profiles of Himalayan bumble bees (Hymenoptera: *Bombus* Latreille) are species-specific and show local and elevation related variation

**DOI:** 10.1101/2023.08.20.554003

**Authors:** Jaya Narah, Martin Streinzer, Jharna Chakravorty, Karsing Megu, Johannes Spaethe, Axel Brockmann, Thomas Schmitt

**Affiliations:** Rajiv Gandhi University, Papum Pare, Arunachal Pradesh, India; National Centre for Biological Sciences - Tata Institute of Fundamental Research, Bengaluru, Karnataka, India; University of Vienna, Vienna, Austria; Chair of Behavioral Physiology and Sociobiology, Biocenter, University of Würzburg, Würzburg, Germany; Department of Animal Ecology and Tropical Biology, Biocenter, University of Würzburg, Würzburg, Germany

**Author notes:** Corresponding author: Thomas Schmitt.

## Abstract

Bumble bees are important pollinators in natural environments and agricultural farmlands and are in particular adapted to harsh environments like high mountain habitats. In these environments, animals are exposed to low temperature and face the risk of desiccation. The Eastern Himalayas are one of the recognized biodiversity hotspots worldwide. The area covers subtropical rainforest with warm temperature and high precipitation as well as high mountain ranges with peaks reaching up to 6000 m, shaping a diverse floral and faunal community at the different altitudinal zones. We investigated the cuticular hydrocarbon profiles of four bumble bee species occurring at different elevational ranges in Arunachal Pradesh, the northeast most state in India. At 17 locations along an elevational gradient we collected workers of two species from lower elevations (*B. albopleuralis* and *B. breviceps;* ∼ 100m - 3000m asl) and two species from higher elevations (*B. prshewalskyi* and *B. mirus*; ∼ 2800m - 4,500m asl). The CHC profiles of all four species showed a significant degree of variation in the composition of hydrocarbons, indicating species specificity. We also found clear correlation with elevation. The weighted mean chain length of the hydrocarbons significantly differed between the low and high altitudinal species, and the proportion of saturated hydrocarbons in CHC profiles significantly increased with the elevational range of the bumble bee species. Thus, these four species of bumble bees in the eastern Himalayas seem to adapt their CHC composition to elevation by decreasing water permeability of their cuticle, similar to insects living in dry mountains or deserts habitats.

## INTRODUCTION

Bumble bees, which evolved in cold temperate environments, occur in India and South Asia only in the mountain ranges of the Himalayas where they play an important role as pollinators of plants and agricultural crops (Rather et al. 2023). The Himalayas are one of the longest mountain ranges in the world and exhibit a steep climatic gradient from the colder and dry western part to the warmer and very humid eastern regions. Corresponding to this gradient, one finds a strong shift in floral composition along the mountain range, from temperate broadleaf forest and arid alpine meadows in the west to wet subtropical broadleaf forest and moist alpine meadows in its east (Rawat 2017). In the eastern part, precipitation can reach up to 5000 mm per year, whereas precipitation decreases with altitude (Dhar and Nandargi 2006). Temperature and rainfall are environmental variables that strongly affect presence and distribution of species. Due to global climate change these local ecosystems might face dramatic changes in the next decades which in turn affect local flora and fauna (Shrestha et al. 2012). For the conservation of Himalayan ecosystems, we need to know the distribution and life histories of plants and animals, and their potential plasticity to respond and adapt to climate changes (Kerr et al. 2015). Due to their importance as pollinators of many flowering plants, bumble bees and other wild bees became a model system to better understand their vulnerability and resilience against climate changes (Soroye et al. 2020; Maebe et al. 2021; Warrit et al. 2023). Possible mechanisms of adaptation include, for example, pilosity (Hines et al. 2022), cuticular hydrocarbons (Maihoff et al. 2023), temperature tolerance (Pimsler et al. 2020; Martinet et al. 2021), respiratory and neural systems (Jackson et al., 2020).

The mountain ranges of the eastern Himalayas in Arunachal Pradesh exhibit one of the largest elevational spans worldwide (from 44 m close to the Bramaputra riverbed up to 7060 m above sea level) and provide a unique opportunity to study bumble bee adaptations to different elevations and associated biotic and abiotic factors by comparing morphological and physiological characters of species or populations with restricted elevation range sizes. Beside the fact that this part of the Himalayas is one of the most biodiverse regions on our planet (Myers et al. 2000), it is also one of the most understudied areas. Thus, a few years ago, we started systematic taxonomic research of the bumble bee fauna in this region (Streinzer et al. 2019).

In the present study we compared the cuticular hydrocarbon (CHC) profiles of two pairs of bumble bee species which either only occur in the lower or higher elevations: *B. albopleuralis* (150 m - 2,990 m), *B. breviceps* (480 m - 2,790 m), *B. prshewalskyi* (4,110 m - 4,400 m), *B. mirus* (2,850 m - 4,260 m) (Streinzer et al. 2019, this publications). It is known that hydrocarbon profiles on the cuticle of insects are highly variable phenotypic character; as the boundary between the organisms and the environment they play a vital role in protecting the body against detrimental abiotic environmental conditions, like heat and desiccation stress, but they also function in intra- and interspecific recognition and communication (Sprenger and Menzel 2020).

The cuticular hydrocarbons can be composed of more than 100 compounds from three substance classes with different physico-chemical properties. Due to these differences CHC profiles can adapt to extreme environmental conditions as drought stress by alternating the composition of compounds with different structural features and chain-length. Reducing the relative number of unsaturated hydrocarbons in favor of n-alkanes will harden the CHC profile and make this hydrophobic waxy layer more waterproof. A similar effect could be achieved by the increase of chain-length of the CHC compounds (Gibbs and Rajpurohit 2010; Menzel et al. 2017). Since we selected bumblebees which either be adapted to low altitude with high humidity and low drought stress or be adapted to high altitude with low humidity and precipitation and high level of drought stress, we hypothesize that high altitude species have a better adapted CHC profile against desiccation than low altitude species. Thus, we expect a higher proportion of saturated hydrocarbons or an increase in the medium chain-length in high altitude species for a better protection against desiccation.

## METHODS AND MATERIALS

### Study area and bumble bee collection

Foraging bumble bee workers were caught at 17 different localities along an elevational transect in the western districts of Arunachal Pradesh between June and September 2021 (Fig. 1). For the current study we selected the CHC profiles of two pairs of low (wet subtropical broadleaf forest) and high altitude (moist alpine meadows) species covering an elevation gradient from 150 to 4,403 m: *Bombus albopleuralis* Friese, 1916 (range: 150 - 1561m, N = 31), *Bombus breviceps* Smith, 1852 (range: 941m - 1575m, N = 12), *B. mirus* (range: 4,136 - 4,403m, N = 29), *B. prshewalskyi:* (range 4,112 - 4,403m, N = 33) (see supplementary Table 1). Collected specimens were brought to the lab and stored at −20 °C before extraction of cuticular hydrocarbons.

**Figure 1.**
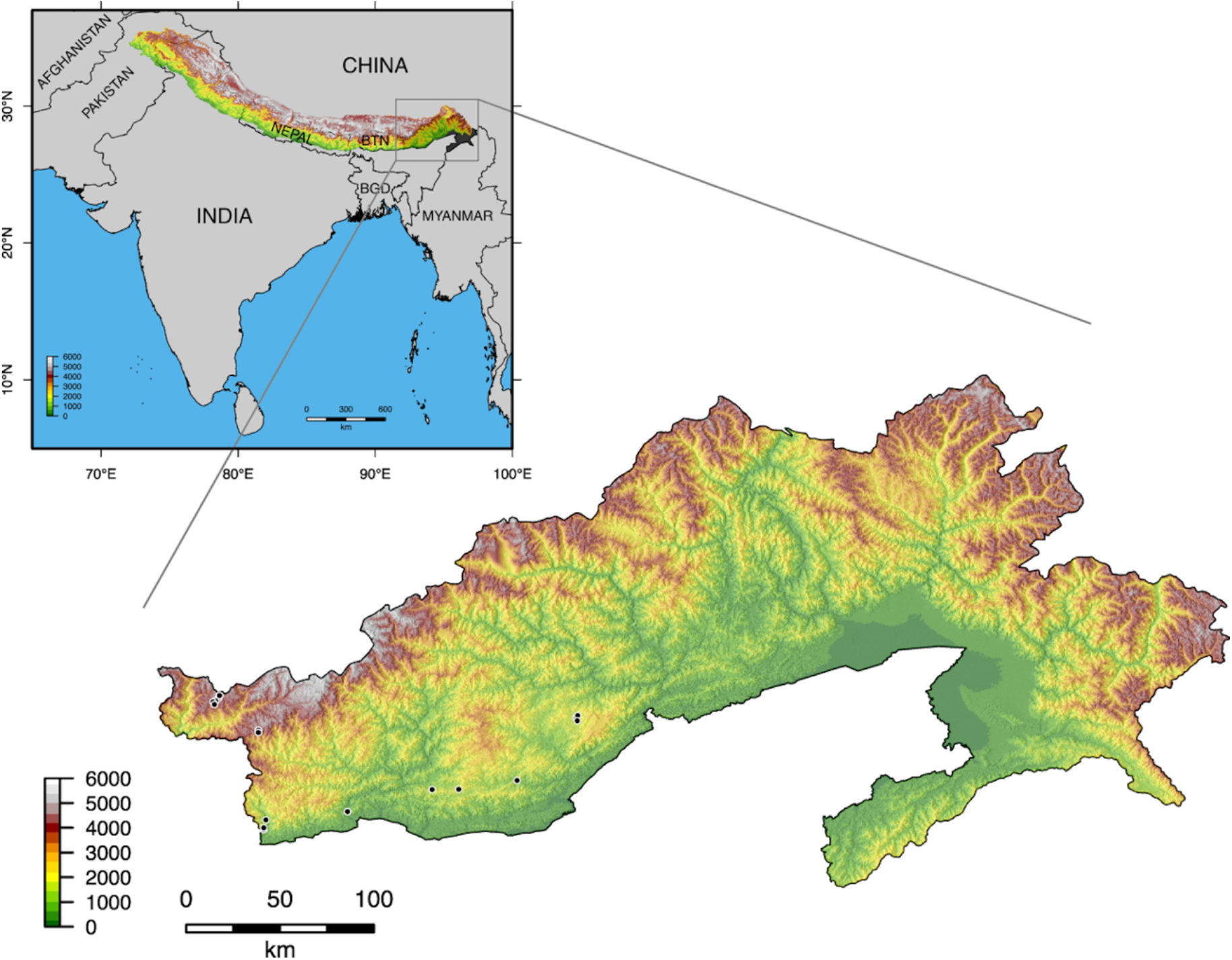
Sampling locations in Arunachal Pradesh. Overview map of the region with an overlay of the elevation profile of the Himalayan Mountain range. Arunachal Pradesh is indicated in darker gray in the overview and shown enlarged at the bottom. Sampling locations (N=17) shown as black dots. Color refers to elevation. Elevation data from (Jarvis et al. 2008), country borders from (Global Administrative Areas 2022) and Himalaya range borders from (Liu and Zhu 2022) (For detailed information about the location and number of collected specimens see ST.1).

### Chemical analyses

CHC profiles were extracted from each worker by immersing the specimen in n-hexane for 10 min. The CHC extracts were stored at −20 C and later reconstituted in approximately 150 µl of hexane for the GC-MS analysis. The extracts were analyzed with an Agilent 7890 gas chromatography coupled with an Agilent 5975 Mass Selective Detector (GC-MS, Agilent, Waldbronn, Germany): The GC (split/splitless injector in splitless mode for 1 min, injected volume 1 µl at 300 °C) was equipped with a DB-5 Fused Silica capillary column (30 m × 0.25 mm ID, df=0.25 µm; J&W Scientific, Folsom, United States). Helium served as carrier gas with a constant flow of 1 mL/min. The following temperature program was used: Start temperature 60 °C, temperature increase by 5 °C per min up to 300 °C, isotherm at 300 °C for 10 min. The electron ionization mass spectra (EI-MS) were acquired at an ionization voltage of 70 eV (source temperature: 230 °C). Chromatograms and mass spectra were recorded with the software HP Enhanced ChemStation G1701AA (version A.03.00; Hewlett Packard). Alongside CHC samples, we run two analytical alkane standards (C8-C20 and C21-C40; Sigma Aldrich) for the calculation of the retention indices. CHC compounds were quantified by integrating peak areas and calculating their relative composition, They were identified by their compound specific retention indices and their diagnostic fragmentation pattern (Carlson et., al. 1998).

### Statistical analysis

All statistical analyses were performed in R, version 4.3.0 (R core team 2023). In all four bumble bee species, we compared the relative abundances of compounds in the CHC profile of workers. CHC compounds were assessed by non-metric multidimensional scaling (NMDS), a two-dimensional ordination method to visualize similarities. Bray-Curtis distance method was used to calculate the dissimilarities between the bumble bee worker profiles. CHC profile composition differences between species were calculated using permutational multivariate analysis of variance (PERMANOVA) in the package vegan (Oksanen et al. 2020) and PairwiseAdonis (Martinez Arbizu 2020) with Bonferroni correction. Permutations were set on 999 or 10,000 respectively.

We determined climate associated chemical traits of the CHC profiles as the proportion of saturated CHCs and abundance of weighted mean chain length. The higher proportion of saturated CHC components and mean chain length enhances the waterproofing in insects cuticle (Gibbs and Pomonis 1995). Differences in weighted mean chain length and proportion of saturated CHCs of the bumble bee workers were analyzed with Kruskal Wallis test and multiple comparisons were performed with Dunn’s test (Dunn 1964, “dunn.test” in R, Dinno 2015).

### COI barcoding and species identification

As species identification based on morphological traits is difficult for bumblebees of the eastern Himalayas, we used DNA barcoding as a more reliable character. DNA was extracted from the muscles of the forelegs or middle legs and isolated using DNeasy Blood & Tissue extraction kit (Qiagen) following the manufacturer’s instructions with slight modifications. (PCR) target region amplification of the standard 658bp barcoding region was performed using standard primers (LepF1: 5’-TTCAACCAATCATAAAGATATTGG-3’ and LepR1: 5’-AACTTCTGGATGTCCAAAAAATCA-3’; Herbert et al. 2004). PCR reactions had a volume of 25µl and contained 0.167 U/µl Taq DNA Polymerase (Invitrogen), 1X Taq Buffer (Invitrogen), 1 mM MgCl2 (Invitrogen), 0.25 mM dNTP mix (Invitrogen), 0.4 µM LepF1, 0.4 µM LepR1, 16.29 µl nuclease free water and 3 µl template DNA. PCR products were verified by gel electrophoresis using 3 µl of PCR product and 2 µl of 100 bp Plus DNA Ladder (Biolabs) and run at 90 V (Tarsons) for 1 hour. The gel was then inspected under UV illumination in a Gel-Doc.

For species verification we used the same amplified DNA extracts. The cytochrome oxidase I gene (COI) was amplified with primer combinations (forward: LepF1 and reverse: LepR1; Herbert et al. 2004) to successfully retrieve overlapping gene fragments merged to a consensus sequence for each specimen. The trace files were manually inspected for errors of the base calling algorithm. Consensus sequences were then exported and aligned in Bioedit (version 7.2 for windows) or Aliview (version 1.2) using MUSCLE algorithm (Edger 2004) and the final alignment was 657 bp long. Primer sequences were removed prior to further analysis.

Reference sequences were selected from Santos Júnior et al. (2022) and downloaded from the BOLD database (Ratnasingham and Herbert 2007), closely related taxa or reliable sequences were picked. If available, sequences from revision studies and designated barcode proxy type specimens were given preference (Williams et al. 2020). In addition, barcode sequences of bumble bees stored at the NCBS research collection facility were used as public reference sequences are not available for species occurring in the eastern Himalayas. The tree was reconstructed using the Maximum Likelihood method using the GTR+GI model (Nei M. and Kumar S. 2000, Kumar S. et al. 2018) for all three codon positions (Kimura 1980) in MEGA (version 10.2 for Windows). Confidence values for nodes were calculated using the bootstrap method with 1000 replicates (Felsenstein 1985). Figtree V1.4.4 was used to view and edit the phylogenetic tree.

## RESULTS

The composition of CHC profiles significantly differed among the four species (PERMANOVA: F= 143.39, df= 3, p<0.001, R^2^ = 0.809) resulting in species specific CHC profiles (Fig. 1; ST: 2 and 3 PERMANOVA between all species pairs: p = 0.001)

The weighted mean chain lengths of hydrocarbons in the CHC profiles significantly differed between species occurring at high and low altitudes (χ2 = 72.165, df = 3, p<0.001, Fig. 3a). The CHC profiles of lower altitude species consisted of hydrocarbons with shorter mean chain length, *Bombus albopleuralis* (mean: 24.9 ± 0.2) and *Bombus breviceps* (mean: 24.5 ± 0.33) whereas the CHC profiles of the higher altitude species contained hydrocarbons with higher weighted mean chain length, *Bombus mirus* (mean: 25.6 ± 0.21) and *Bombus prshewalskyi* (mean: 25.5 ± 0.26).

**Figure 2.**
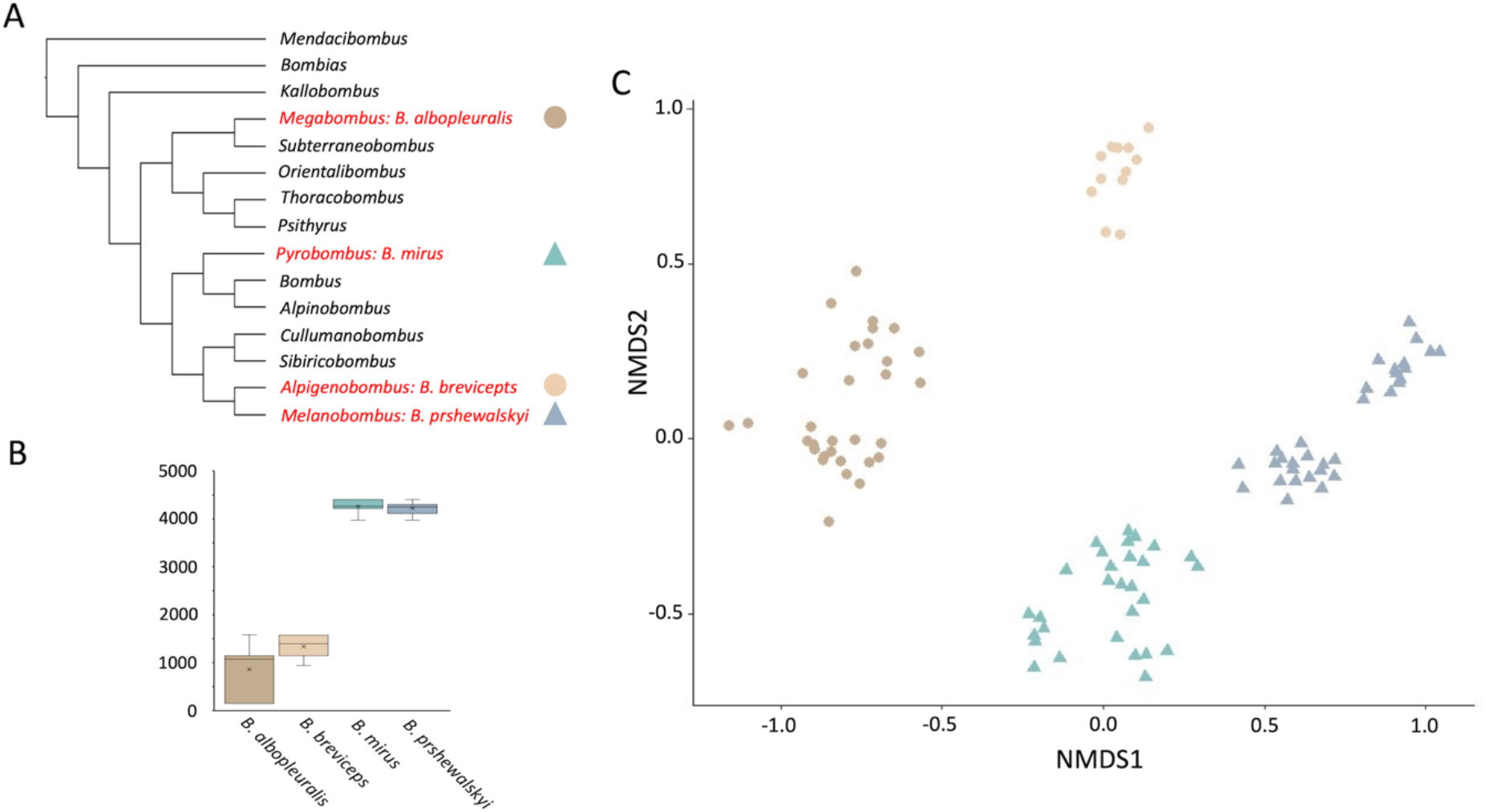
CHC profiles diversity of four Himalayan bumble bee species with different elevational ranges. (A). Phylogenetic position of the study species embedded in the subgeneric classification of the bumble bees (Sun et al., 2021, Williams et al., 2022). (B.) Elevational range of collected bumble bee specimens. (C.) Diversity of CHC profiles of bumble bees displayed in a two-dimensional graph by non-metric multidimensional scaling (NMDS) based on Bray-Curtis distances. Distance between symbols indicates the degree of similarity among the CHC profiles. Each symbol represents the CHC profile of an individual bumble bee worker. Circles: species with lower elevational ranges, triangles: species with lower elevational ranges. Colors indicate the species.

**Figure 3.**
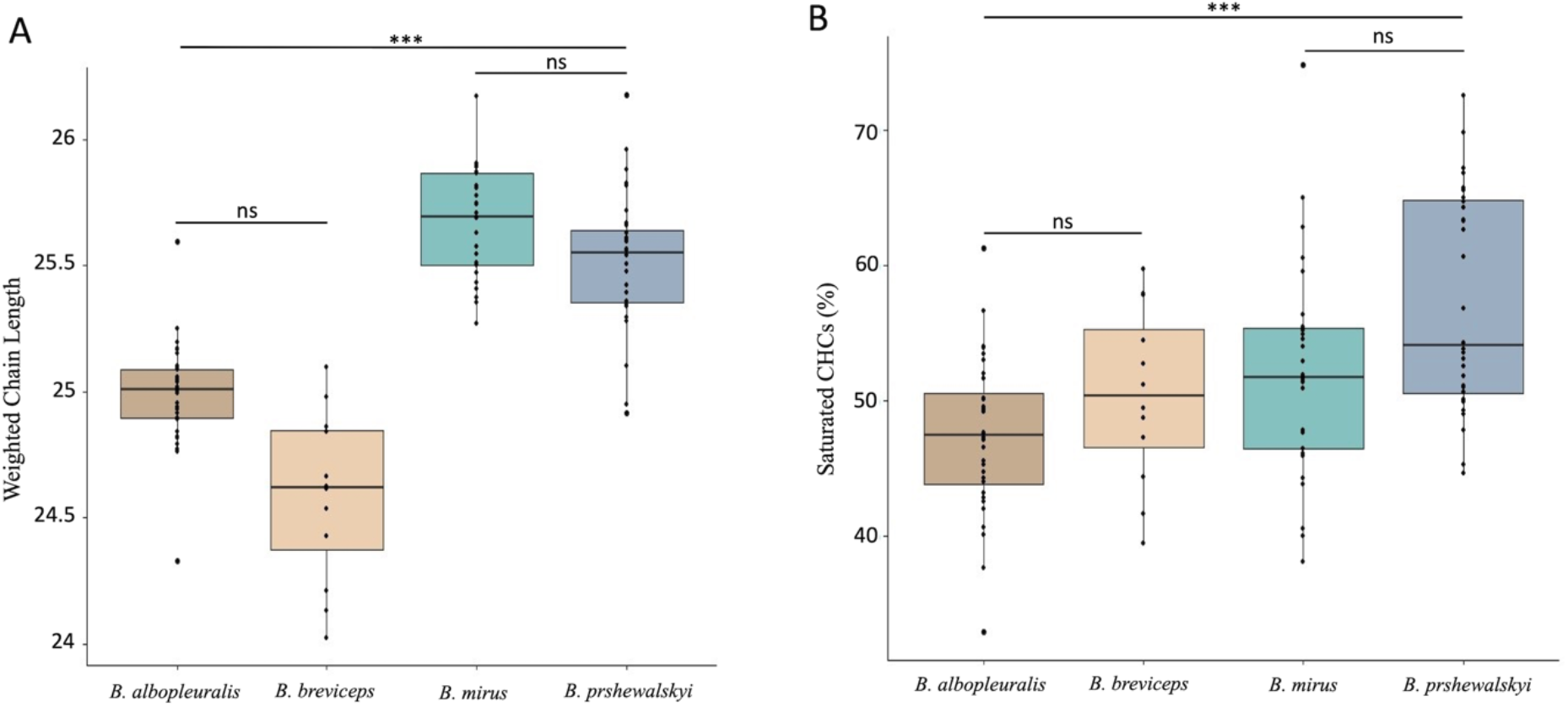
Differences in climate associated chemical traits of CHC profiles in the four bumble bee species. (A) Differences in weighted mean chain lengths of the hydrocarbons in the CHC profiles (Kruskal Wallis test and multiple comparisons using Dunn’s test (Dunn 1964, “dunn.test” in R, Dinno 2015). (B) Differences in the proportion of saturated hydrocarbons in the CHC profiles. Kruskal Wallis test and multiple comparisons using Dunn’s test (Dunn 1964, “dunn.test” in R, Dinno 2015)

The proportion of saturated CHC components in the CHC profile increased with the elevational range of the bumble species (χ2 = 2.521, df = 3, p<0.001) (Fig. 3b). *Bombus albopleuralis*, the species with the lowest elevational range, had the lowest mean proportion of saturated CHC components (average: 47% ± 5.7%) followed by *Bombus breviceps* (average: 50% ± 6.4%), and the two high elevation species, *Bombus mirus* (average: 52% ± 7.8%) and *Bombus prshewalskyi* (average: 57% ± 7.9%).

## DISCUSSION

In this study, we explored the variation in CHC profiles of four bumble bee species with different elevational ranges to identify possible adaptations to changes in climate and desiccation stress with altitude. Bumble bees often occur in a wide altitudinal range, but only a few species occur in tropical lowlands, where the conditions are typically not suitable for these bees that are adapted to cold environments (Moure and Sakagami 1962; Williams 1991; Gonzalez et al. 2004; Williams et al. 2009). In the mountain regions of Arunachal Pradesh, the highest bumble bee diversity occurs in the subalpine alpine zones at the higher altitudes and it decreases towards the lower elevations (Streinzer et al. 2019). Bumble bee species preferring high or low altitudinal niches revealed distinct and species-specific CHC profiles. These differences are particularly pronounced by the presence of methyl-branched alkanes in high altitude species, whereas these compounds are absent in both low altitude species. In addition to species-specificity of CHC profiles, the NMDS also revealed subclusters for all species except *B. breviceps*. Since these subclusters are composed of specimens collected at different locations they likely represent different populations. In this case, the CHC profile subcluster might be the result of neutral evolution due to reduced or absent gene flow between these allopatrically isolated populations.

Compositional features in CHC profiles are explained by similarities in the elevational niche preferences of the species, indicating that CHC profiles found on the cuticle of these species are shaped by environmental factors correlating with elevation (e.g., temperature, precipitation, humidity). High altitudes are often drier, with less precipitation, and less humid than lower altitudes. This is particularly true for the Eastern range of the Himalaya (Dhar and Nandargi 2006). To cope with drought stress insects harden their CHC profile as a plastic change of their composition (Stinziano et al 2015) or as a genetically fixed adaptation (Rajpurohit et al. 2017). CHC profiles can either be hardened by elongating the chain-length of their compounds or change the relative composition to a higher proportion of saturated against unsaturated hydrocarbons. In our study we find both features realized by the investigated bumblebees. The weighted chain-length is elongated in the high-altitude species compared to both low altitude species. The variation of the ratio of saturated to unsaturated hydrocarbons is less pronounced. However, our study reveals a significant increase in the ratio by comparing *B. alpopleuralis* as a low altitude species with *B. prshewalskyi* as a high-altitude species.

The results of our study are congruent with a study of halictid bees on the slope of the Kilimanjaro. Bees of the genus Lasioglossum changed the composition of their CHC profiles similar to the bumblebees investigated in the present study (Mayr et al. 2021). In contrast, a study on bumble bees from an elevational gradient in the Alps did not reveal an adaptation of the CHC profiles to potential drought stress in higher altitudes (Maihoff et al. 2023). The elevational gradient of 1000 m studied might have been too small to cause pronounced changes in the environmental conditions and thus, the selection pressure might not be strong enough to change the compositional features of the CHC profiles in the selected bumble bee species. Although we do not provide bioassays testing actual desiccation stress, we know from other studies that hardening of a CHC profile is a direct reaction to desiccation stress (Stinziano et al. 2015; Rajpurohit et al. 2017) and can thus be suggested as adaptation to the harsher environment on high altitudes in the eastern Himalaya.

## ACKNOWLEDGEMENTS

We are grateful to B. Narah who help with the collection of bumble bee specimens and A. Suryanarayan and Prabhu MV for help with the identification of CHC profiles and statistical analysis. We thank Dr. Yeshwanth HM and NCBS Research Collection for constant support and help during the study. The study is part of the Chemical Ecology Network Programme funded by the Department of Biotechnology, Govt of India (DBT-NER/Agri/24/2013). The field trips of researchers from RGU and NCBS were supported by funds from the Chemical Ecology Project as well as NCBS/TIFR institutional funds [12P4167] to AB. We acknowledge support of the Department of Atomic Energy, Government of India [under project no. 12-R&D-TFR-5.04-0800]. Permission to collect bumble bees was granted by the Arunachal Pradesh Biodiversity Board (SFRI/APBB/09/2022/581071). Researchers from the University of Würzburg and Vienna were supported by institutional funds from the University of Würzburg to Thomas Schmitt and Johannes Spaethe.

## SUPPLEMENTARY MATERIAL

**Supplementary Table 1:** List of collected worker specimens.

| Elevation (m) | Species | Number | Locality |
| --- | --- | --- | --- |
| 150 | *B. albopleuralis* | 13 | N27°4.215', E092°35.472'<br>Bhalukpong, West Kameng |
| 800 | *B. albopleuralis* | 10 | N26°58.290', E092°7.972'<br>Balemu, West Kameng |
| 941 | *B. breviceps* | 2 | N27°14.465', E093°31.144'<br>Toru, Sagalee, Papum Pare |
| 1145 | *B. albopleuralis* | 5 | N27°11.462', E093°3.261'<br>Papu Valley, West Kameng |
|  | *B. breviceps* | 2 |  |
| 1217 | *B. breviceps* | 2 | N27°1.482', E092°8.674'<br>Ankalin, West Kameng |
| 1445 | *B. albopleuralis* | 1 | N27°11.542', E093°11.997'<br>10 km ahead Pakke Kessang, |
| 1561 | *B. albopleuralis* | 2 | N27°35.126', E093°50.598'<br>Kalung Village, Ziro Valley |
|  | *B. breviceps* | 2 |  |
| 1575 | *B. breviceps* | 4 | N27°35.817', E093°51.47'<br>Salaya, Ziro Valley |
| 4112 | *B. prshewalskyi* | 12 | N27°31.011', E092°6.203'<br>Sela Pass, West Kameng |
| 4136 | *B. mirus* | 5 | N27°39.478', E091°51.778'<br>Nagula, Tawang |
| 4215 | *B. prshewalskyi* | 2 | N27°30.228', E092°6.205'<br>Sela top, West Kameng |
| 4247 | *B. prshewalskyi* | 7 | N27°40.666', E091°52.202'<br>Klemta Check gate, Tawang |
|  | *B. mirus* | 9 |  |
| 4264 | *B. prshewalskyi* | 2 | N27°40.299', E091°51.339'<br>10km N Tawang, Tawang |
|  | *B. mirus* | 7 |  |
| 4300 | *B. prshewalskyi* | 10 | N27°41.745', E91°53.260'<br>Klemta, Tawang |
|  | *B. mirus* | 1 |  |
| 4403 | *B. mirus* | 7 | N27°42.412', E091°53.455'<br>Bumla Pass, Tawang |

**Supplementary Table 2.**
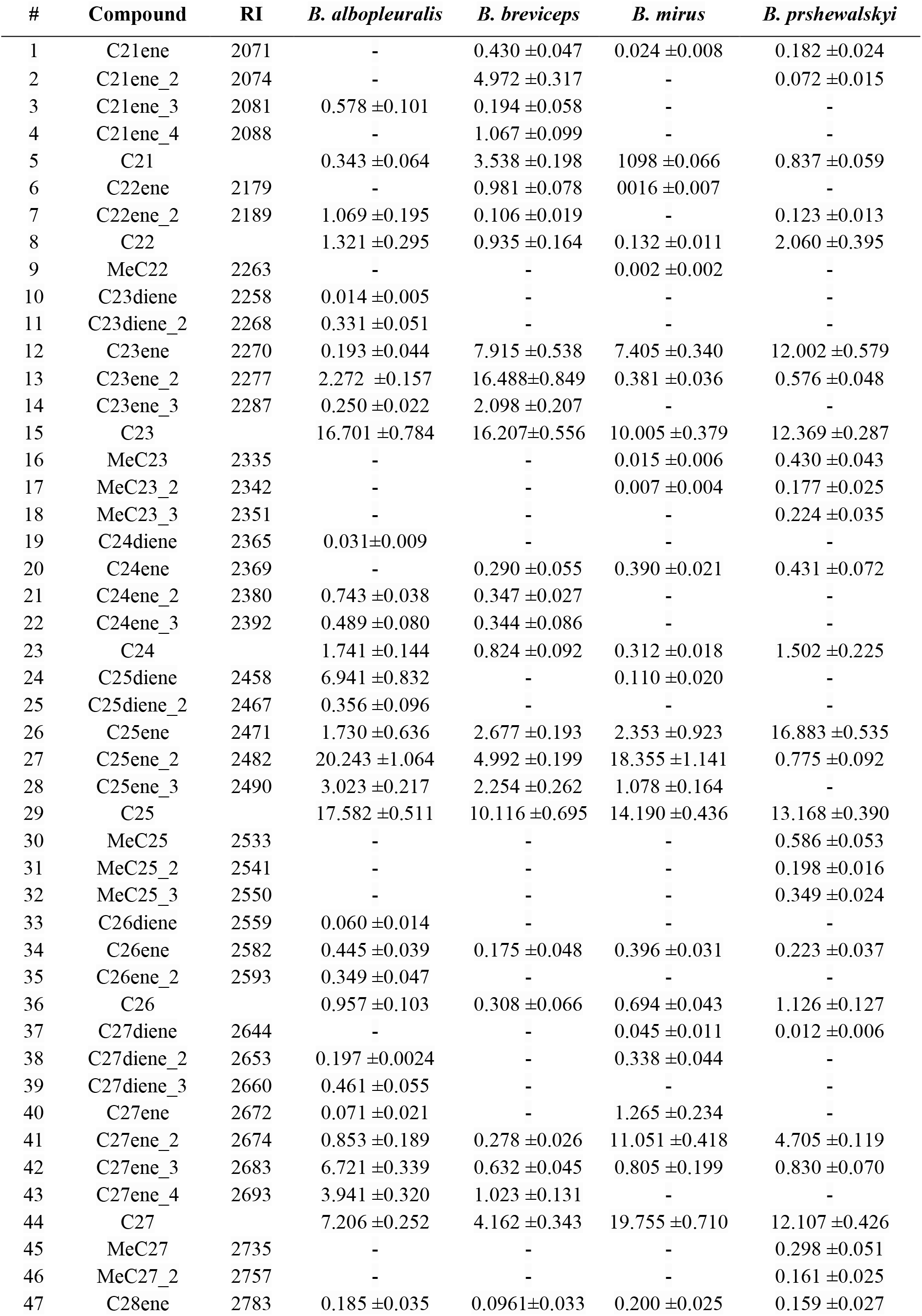
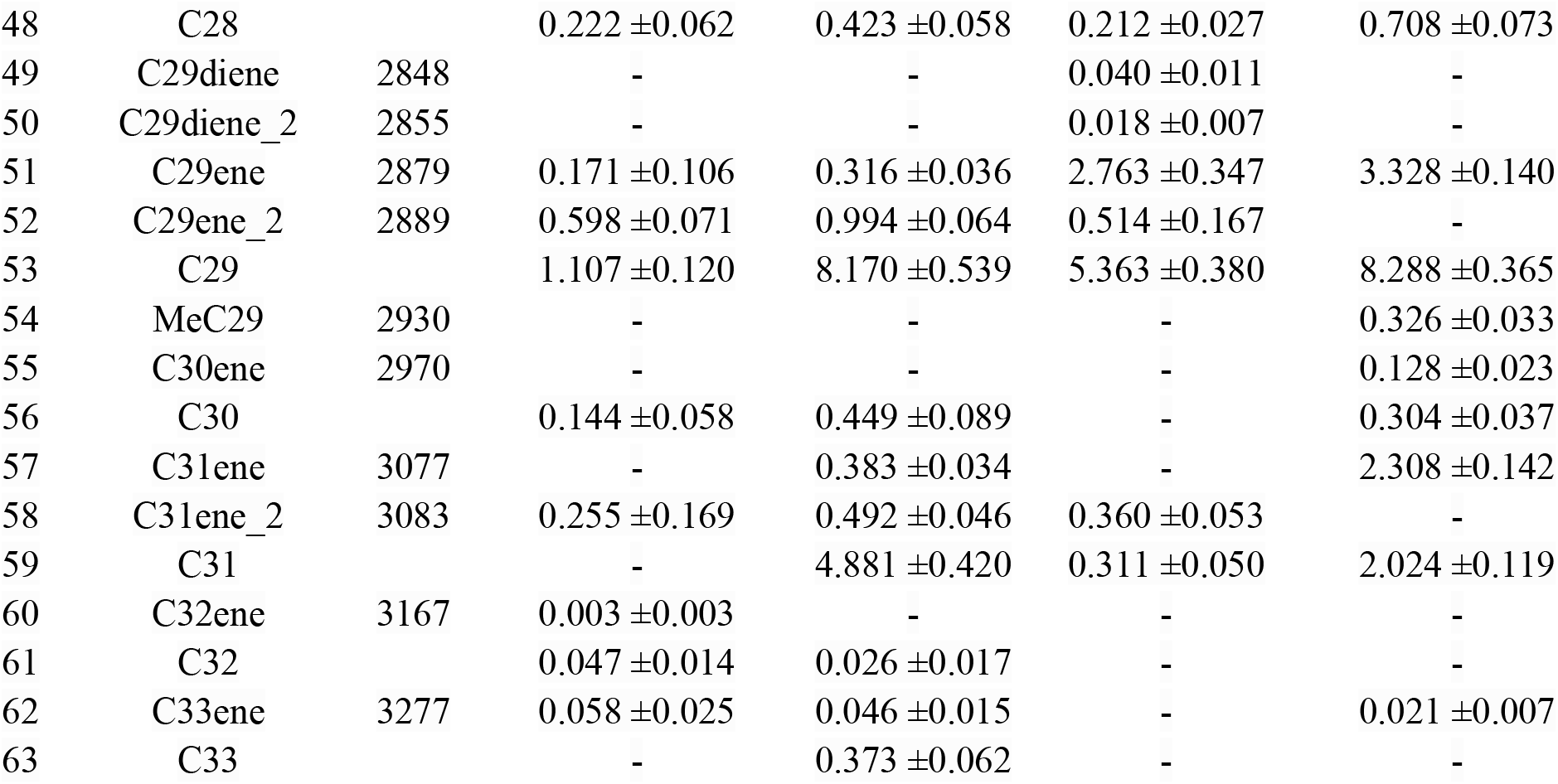
List of peaks of cuticular hydrocarbons from low and high bumble bee species.

| # | Compound | RI | *B. albopleuralis* | *B. breviceps* | *B. mirus* | *B. prshewalskyi* |
| --- | --- | --- | --- | --- | --- | --- |
| 1 | C21ene | 2071 | - | 0.430 ±0.047 | 0.024 ±0.008 | 0.182 ±0.024 |
| 2 | C21ene_2 | 2074 | - | 4.972 ±0.317 | - | 0.072 ±0.015 |
| 3 | C21ene_3 | 2081 | 0.578 ±0.101 | 0.194 ±0.058 | - | - |
| 4 | C21ene_4 | 2088 | - | 1.067 ±0.099 | - | - |
| 5 | C21 |  | 0.343 ±0.064 | 3.538 ±0.198 | 1098 ±0.066 | 0.837 ±0.059 |
| 6 | C22ene | 2179 | - | 0.981 ±0.078 | 0016 ±0.007 | - |
| 7 | C22ene_2 | 2189 | 1.069 ±0.195 | 0.106 ±0.019 | - | 0.123 ±0.013 |
| 8 | C22 |  | 1.321 ±0.295 | 0.935 ±0.164 | 0.132 ±0.011 | 2.060 ±0.395 |
| 9 | MeC22 | 2263 | - | - | 0.002 ±0.002 | - |
| 10 | C23diene | 2258 | 0.014 ±0.005 | - | - | - |
| 11 | C23diene_2 | 2268 | 0.331 ±0.051 | - | - | - |
| 12 | C23ene | 2270 | 0.193 ±0.044 | 7.915 ±0.538 | 7.405 ±0.340 | 12.002 ±0.579 |
| 13 | C23ene_2 | 2277 | 2.272 ±0.157 | 16.488±0.849 | 0.381 ±0.036 | 0.576 ±0.048 |
| 14 | C23ene_3 | 2287 | 0.250 ±0.022 | 2.098 ±0.207 | - | - |
| 15 | C23 |  | 16.701 ±0.784 | 16.207±0.556 | 10.005 ±0.379 | 12.369 ±0.287 |
| 16 | MeC23 | 2335 | - | - | 0.015 ±0.006 | 0.430 ±0.043 |
| 17 | MeC23_2 | 2342 | - | - | 0.007 ±0.004 | 0.177 ±0.025 |
| 18 | MeC23_3 | 2351 | - | - | - | 0.224 ±0.035 |
| 19 | C24diene | 2365 | 0.031±0.009 | - | - | - |
| 20 | C24ene | 2369 | - | 0.290 ±0.055 | 0.390 ±0.021 | 0.431 ±0.072 |
| 21 | C24ene_2 | 2380 | 0.743 ±0.038 | 0.347 ±0.027 | - | - |
| 22 | C24ene_3 | 2392 | 0.489 ±0.080 | 0.344 ±0.086 | - | - |
| 23 | C24 |  | 1.741 ±0.144 | 0.824 ±0.092 | 0.312 ±0.018 | 1.502 ±0.225 |
| 24 | C25diene | 2458 | 6.941 ±0.832 | - | 0.110 ±0.020 | - |
| 25 | C25diene_2 | 2467 | 0.356 ±0.096 | - | - | - |
| 26 | C25ene | 2471 | 1.730 ±0.636 | 2.677 ±0.193 | 2.353 ±0.923 | 16.883 ±0.535 |
| 27 | C25ene_2 | 2482 | 20.243 ±1.064 | 4.992 ±0.199 | 18.355 ±1.141 | 0.775 ±0.092 |
| 28 | C25ene_3 | 2490 | 3.023 ±0.217 | 2.254 ±0.262 | 1.078 ±0.164 | - |
| 29 | C25 |  | 17.582 ±0.511 | 10.116 ±0.695 | 14.190 ±0.436 | 13.168 ±0.390 |
| 30 | MeC25 | 2533 | - | - | - | 0.586 ±0.053 |
| 31 | MeC25_2 | 2541 | - | - | - | 0.198 ±0.016 |
| 32 | MeC25_3 | 2550 | - | - | - | 0.349 ±0.024 |
| 33 | C26diene | 2559 | 0.060 ±0.014 | - | - | - |
| 34 | C26ene | 2582 | 0.445 ±0.039 | 0.175 ±0.048 | 0.396 ±0.031 | 0.223 ±0.037 |
| 35 | C26ene_2 | 2593 | 0.349 ±0.047 | - | - | - |
| 36 | C26 |  | 0.957 ±0.103 | 0.308 ±0.066 | 0.694 ±0.043 | 1.126 ±0.127 |
| 37 | C27diene | 2644 | - | - | 0.045 ±0.011 | 0.012 ±0.006 |
| 38 | C27diene_2 | 2653 | 0.197 ±0.0024 | - | 0.338 ±0.044 | - |
| 39 | C27diene_3 | 2660 | 0.461 ±0.055 | - | - | - |
| 40 | C27ene | 2672 | 0.071 ±0.021 | - | 1.265 ±0.234 | - |
| 41 | C27ene_2 | 2674 | 0.853 ±0.189 | 0.278 ±0.026 | 11.051 ±0.418 | 4.705 ±0.119 |
| 42 | C27ene_3 | 2683 | 6.721 ±0.339 | 0.632 ±0.045 | 0.805 ±0.199 | 0.830 ±0.070 |
| 43 | C27ene_4 | 2693 | 3.941 ±0.320 | 1.023 ±0.131 | - | - |
| 44 | C27 |  | 7.206 ±0.252 | 4.162 ±0.343 | 19.755 ±0.710 | 12.107 ±0.426 |
| 45 | MeC27 | 2735 | - | - | - | 0.298 ±0.051 |
| 46 | MeC27_2 | 2757 | - | - | - | 0.161 ±0.025 |
| 47 | C28ene | 2783 | 0.185 ±0.035 | 0.0961±0.033 | 0.200 ±0.025 | 0.159 ±0.027 |
| 48 | C28 |  | 0.222 ±0.062 | 0.423 ±0.058 | 0.212 ±0.027 | 0.708 ±0.073 |
| 49 | C29diene | 2848 | - | - | 0.040 ±0.011 | - |
| 50 | C29diene_2 | 2855 | - | - | 0.018 ±0.007 | - |
| 51 | C29ene | 2879 | 0.171 ±0.106 | 0.316 ±0.036 | 2.763 ±0.347 | 3.328 ±0.140 |
| 52 | C29ene_2 | 2889 | 0.598 ±0.071 | 0.994 ±0.064 | 0.514 ±0.167 | - |
| 53 | C29 |  | 1.107 ±0.120 | 8.170 ±0.539 | 5.363 ±0.380 | 8.288 ±0.365 |
| 54 | MeC29 | 2930 | - | - | - | 0.326 ±0.033 |
| 55 | C30ene | 2970 | - | - | - | 0.128 ±0.023 |
| 56 | C30 |  | 0.144 ±0.058 | 0.449 ±0.089 | - | 0.304 ±0.037 |
| 57 | C31ene | 3077 | - | 0.383 ±0.034 | - | 2.308 ±0.142 |
| 58 | C31ene_2 | 3083 | 0.255 ±0.169 | 0.492 ±0.046 | 0.360 ±0.053 | - |
| 59 | C31 |  | - | 4.881 ±0.420 | 0.311 ±0.050 | 2.024 ±0.119 |
| 60 | C32ene | 3167 | 0.003 ±0.003 | - | - | - |
| 61 | C32 |  | 0.047 ±0.014 | 0.026 ±0.017 | - | - |
| 62 | C33ene | 3277 | 0.058 ±0.025 | 0.046 ±0.015 | - | 0.021 ±0.007 |
| 63 | C33 |  | - | 0.373 ±0.062 | - | - |

**Supplementary Table 3:** Pairwise PERMANOVA to compare differences in CHC profile composition between species.

| Pairs | Df | Sums of Sqs | F-Model | R<sup>2</sup> | p-value |
| --- | --- | --- | --- | --- | --- |
| *B. albopleurialis* vs *B. breviceps* | 1 | 2.240 | 97.676 | 0.699 | 0.001 |
| *B. albopleurialis* vs *B. mirus* | 1 | 2.542 | 108.37 | 0.647 | 0.001 |
| *B. albopleurialis* vs *B. prshewalskyi* | 1 | 4.893 | 239.35 | 0.794 | 0.001 |
| *B. breviceps* vs *B. mirus* | 1 | 2.155 | 128.43 | 0.767 | 0.001 |
| *B. breviceps* vs *B. prshewalskyi* | 1 | 1.812 | 141.54 | 0.771 | 0.001 |
| *B. mirus* vs *B. prshewalskyi* | 1 | 2.043 | 125.71 | 0.681 | 0.001 |

